# Chromosomal resistance mutations facilitate acquisition of multidrug-resistance plasmids in *Escherichia coli*

**DOI:** 10.1101/2025.04.15.648949

**Authors:** Khadija-Siddiqa N. Hanga, Michael A. Brockhurst, Michael J. Bottery

## Abstract

Bacteria can gain multiple resistance mechanisms in a single step by the acquisition of multidrug-resistance (MDR) plasmids, but it is unclear how antibiotic selection during the acquisition of MDR plasmids affects the evolution of additional resistance mechanisms. Through conjugating separate ESBL- and carbapenamase-producing MDR plasmids into plasmid-naive *Escherichia coli* hosts we examine the effects of acquisition of a single plasmid or co-acquisition of multiple plasmids upon fitness costs, resistance and subsequent genomic adaptation. We show that acquisition of pOXA-48, encoding OXA-48 carbapenamase, is associated with highly variable fitness costs and levels of resistance to ertapenem in transconjugants independent of the presence of pLL35. This phenomenon was not observed during the acquisition of ESBL CTX-M-15 encoding pLL35 alone. Transconjugants receiving pOXA-48 rapidly gained parallel mutations affecting the membrane porin OmpF, or its regulators OmpR or EnvZ. These chromosomal mutations were not compensatory for the fitness costs imposed by the plasmid, nor did they provide significant increases in resistance to carbapenems in the absence of the pOXA-48. Rather, they acted synergistically with the plasmid-encoded carbapenamase, which alone only provided marginal resistance, together providing high-level resistance to ertapenem. Such rapid evolutionary processes may play an important role in plasmid dynamics within environments with strong antibiotic selection for plasmid-encoded ARGs, particularly when these ARGs provide only marginal resistance.

## Introduction

Antibiotic resistance is a growing public health crisis, with an estimated 1.27 million deaths attributed to antibiotic-resistant infections in 2019 alone [1]. Resistance mechanisms in bacteria can be intrinsic, due to inherent structural and functional traits, or acquired through mutations or horizontal gene transfer (HGT) of antimicrobial resistance genes (ARGs) encoded on mobile genetic elements (MGEs) [2, 3]. MGEs can promote the accumulation and rapid spread of multiple resistance mechanisms in a single step. When bacteria acquire MGEs encoding ARGs, antibiotics may become ineffective, allowing strains to grow in the presence of otherwise lethal antibiotic concentrations and pose a significant challenge to treatment [3].

In recent decades, the rise of carbapenem-resistant Enterobacteriaceae (CRE) and extended-spectrum β-lactamase (ESBL) producing Enterobacteriaceae has been particularly concerning. Infections caused by these antibiotic-resistant bacteria, especially in intensive care units (ICUs) and among patients with severe underlying conditions, are associated with high rates of morbidity and mortality [2, 4, 5]. Resistance within CRE is primarily spread through HGT mediated by conjugative plasmids encoding carbapenem-hydrolysing carbapenemase enzymes [6–9]. Other mechanisms, such as porin mutations and efflux pump overexpression, can also confer resistance to carbapenems, particularly when accompanying β-lactamase enzymes [5, 10].

Although Enterobacteriaceae, such as *Escherichia coli* and *Klebsiella pneumoniae*, form part of the healthy human gut flora, they are also frequently associated with human and animal infections. Invasive illnesses caused by multidrug-resistant (MDR) lineages of Enterobacteriaceae pose a significant barrier to successful treatment. Such invasive disease-causing lineages are often associated with the acquisition of one or more MDR plasmids [11]. Plasmids carried by MDR strains frequently encode ESBLs, and due to the increase in the usage of carbapenems to treat infections, the co-occurrence of separate carbapenem resistance plasmids is on the rise [3, 11]. Combatting such increasing rates of MDR Enterobacteriaceae infections requires understanding of how the carriage of multiple MDR plasmids affects bacterial fitness and resistance levels.

While acquiring an MDR plasmid can provide significant fitness advantages during antibiotic treatment, plasmids often impose a fitness cost in the absence of antibiotics. Such fitness costs can arise through multiple different and often interacting mechanisms, including dysregulation of metabolic pathways, producing and accumulating toxic products, and altering gene expression [12–14]. The carriage of multiple plasmids can have complex effects on fitness costs. Positive epistasis between plasmids costs, whereby the cost of acquiring additional plasmids is less than the sum of the additive costs imposed by each individual plasmid, can promote the accumulation of multiple plasmids within single cells [13, 15], however positive epistasis is not universally observed [16]. The balance between the cost imposed by an MDR plasmid in the absence of antibiotic selection and the benefit in its presence is a critical factor in the persistence of resistance plasmids within bacterial populations [13, 17]. However, fitness costs associated with plasmids can be ameliorated by compensatory mutations either on the chromosome, plasmid or both, stabilising plasmids within bacterial lineages [4, 18–20] and can occur extremely rapidly, even during the outgrowth of transconjugant colonies [21]. Moreover, several experimental studies report that single compensatory mutations can ameliorate the costs of multiple distinct plasmids [16, 20], favouring the maintenance of multiple plasmids within lineages.

Here, we examine the effects of coinfection by two clinical conjugative plasmids, pLL35 and pOXA-48, on resistance, fitness costs, and compensatory mutations. The 106 kb, multidrug-resistant, IncFII(K)-9 group plasmid pLL35 was originally isolated from *K. pneumoniae* and encodes the ESBL CTX-M-15 conferring resistance to cefotaxime which is the most commonly occurring ESBL worldwide [22, 23]. pLL35 is highly stable due to its toxin/antitoxin stability systems [16, 24]. The 65 kb IncL group pOXA-48 plasmid was isolated from a separate *K. pneumoniae* infection and encodes the carbapenemase OXA-48, which accounts for 52% of carbapenem resistance cases referred to the UK AMRHAI Reference Unit in 2018 [14, 25]. We measured resistance to cefotaxime (CEF) or ertapenem (ERT) and growth kinetics without antibiotics for *E. coli* MG1655 either singly or coinfected with pLL35 and pOXA-48 and obtained whole genome sequences for multiple independent transconjugants. We found that acquisition of pOXA-48, either alone or together with pLL35, was facilitated by chromosomal resistance mutations affecting genes encoding outer membrane proteins. Using Keio single gene knockout strains, we show that the mutated chromosomal loci are not classical compensatory mutations but instead act synergistically with the plasmid’s OXA-48 ARG to provide high-level resistance to ertapenem.

## Methods

### Bacterial strains, plasmids and culture conditions

Two isogenic strains of *E. coli* K12 MG1655 chromosomally labelled with EGFP (MG1655::GFP; subsequently denoted MG1655G) or mCherry (MG1655::mCherry subsequently denoted MG1655R) at the attB natural insertion site through λ red homologous recombination were used as recipients for conjugation experiments unless otherwise stated [19]. Fluorescently labelled *E. coli* MG1655 strains were also labelled with kanamycin resistance (NeoR). The plasmid donors used for conjugations were *E. coli* J53 for pOXA-48 and *E. coli* MG1655 for pLL35, both established by conjugation from their natural *K. pneumoniae* hosts (Table S1).

All strains were cultured in liquid Nutrient Broth (NB) (Oxoid) at 37°C, shaken at 180 r.p.m, in 6 ml media in 50 ml falcon tubes or on Nutrient Agar (Oxoid) for culture on solid media. Resistance profiles of all strains were confirmed by plating onto solid media containing kanamycin (25 µg/ml) for fluorescent recipients, sodium azide (100 µg/ml) for *E. coli* J53, CEF (4 µg/ml) for strains containing pLL35 or ERT (0.5 µg/ml) for strains containing pOXA-48.

### Plasmid conjugation

Twenty-four independent conjugation assays were conducted to construct eight biological replicates of MG1655 pLL35, MG1655 pOXA-48 and MG1655 pLL35 + pOXA-48 transconjugants. The pLL35 and pOXA-48 resistance plasmids were transferred individually into *E. coli* MG1655 via conjugation. Overnight cultures of recipients (MG1655G and MG1655R) and donors (MG1655 pLL35 and J53 pOXA-48) were inoculated in 5ml of nutrient broth from single colonies isolated from selective plates. pLL35 was conjugated into donors through liquid conjugation, donor and recipient overnight cultures were mixed at ratios of 1:3 for pLL35 and incubated at 37°C in non-selective NB without shaking for 24h. pOXA-48 conjugation was conducted on solid agar, donor and recipients were mixed 1:3 and 50 µl of the mixture was added on sterile filter papers placed on non-selective nutrient agar plates and incubated at 37°C for 24h. To create the double plasmid bearing *E. coli* MG1655 pLL35+pOXA-48 a second round of conjugation was performed using fluorescent *E. coli* MG1655::GFP pOXA-48 and *E. coli* MG1655::mCherry pLL35 at a 1:1 ratio. Transconjugants were selected by plating serial dilutions of the mixed cultures onto nutrient agar supplemented with either CEF, ERT or both. All transconjugants were validated using a multiplex PCR assay designed to amplify the backbone of the two MDR plasmids and their resistance genes separately (Table S2). All subsequent experiments were conducted using six independent validated transconjugants.

### Growth curves

Growth curves were conducted to evaluate the growth kinetics of the newly constructed plasmid-bearing strains and the isogenic parental plasmid free strain. Each transconjugant was individually streaked on the appropriate antibiotic selection plates and incubated overnight at 37°C. The plasmid-free parental strains were also cultured on nutrient agar plates. Three individual replicate colonies were picked from each growth plate with sterile cocktail sticks into 96 well plates containing 200µl of NB, which were incubated overnight at 37°C. The saturated overnight cultures were diluted to an optical density (OD600) of 0.05 in fresh NB in a 96-well plate. This was incubated at 37°C with shaking at 180rpm, 3mm orbital radius for 24hrs in a Tecan Infinite M200 plate reader. The OD600 was measured every 20 minutes and growth parameters of lag time, maximum growth rate, and maximum OD600 and area under the curve were determined using “ggplot2” and “dplyr” packages in Rstudio 2022.07.2.

### Minimum Inhibitory Concentration (MIC) assay

Minimum inhibitory concentration (MIC) assays for cefotaxime and ertapenem were performed using the broth microdilution method described by Wiegand et al. [26]. Single colonies of the parental plasmid-free strain and the pLL35, pOXA-48, and pLL35+pOXA-48 harbouring transconjugants were picked into 5 mL of NB broth from selective agar plates grown overnight at 37°C with shaking at 180 rpm. Overnight cultures were diluted to 5 × 10^6^ CFU/mL in fresh NB. These were then used to inoculate 96-well plates at a final density of 5 × 10^5^ CFU/mL, containing a 2-fold dilution series of CEF, 1 µg/mL to 128 µg/mL, or ERT,0.05 µg/mL to 24 µg/mL. Inoculated 96 plates were incubated at 37°C for 24 hours with no shaking, after which OD600 readings were recorded to assess bacterial growth. The MIC was defined as the lowest concentration of antibiotic that completely inhibits visible growth [26] which is less than 10% of the no-antibiotic control. The clinical MIC breakpoints used for interpretation were 0.5 mg/L for ERT and 4 mg/L for CEF, as per EUCAST guidelines [27].

### Whole Genome Sequencing

Whole genome sequencing was employed to identify mutations in each of the transconjugant replicate strains relative to the parental MG1655 strains. DNA extraction and libraries were prepared for short-read sequencing by MicrobesNG (http://www.microbesng.com), and sequencing was conducted using an Illumina NovaSeq 6000, generating 2 × 250 bp paired-end reads. Variants were identified with Breseq [28] using the *E. coli* MG1655 genome (GenBank accession U00096.3), pOXA-48 (GenBank accession number MT441554) and pLL35 [24] sequences as a reference. Variants identified within the parental plasmid-free *E. coli* MG1655 strain, which were also present within the transconjugants were removed from the analysis to ensure that only newly acquired mutations were annotated.

### Functional analysis using Gene Knockouts (Keio collection)

Gene deletion strains from the Keio collection [29] were used to investigate the impact of loss of function mutations observed within genes identified to have gained mutations during acquisition of either plasmid. Parallel mutations within *ompF, ompR*, and *envZ* were observed upon acquisition of pOXA-48 thus, pOXA-48 was conjugated into *E. coli* BW25113 *ΔompF, E. coli* BW25113 *ΔompR*, and *E. coli* BW25113 *ΔenvZ* together with the parental *E. coli* BW25113 strain. Growth curves of all plasmid-free and pOXA-48-bearing deletion strains and the parental strain were conducted as described above in NB over 24 hours at 37°C. The resistance profiles of the Keio transconjugants to ERT were also obtained using the MIC protocols described above.

## Results

### Effects of MDR plasmids on resistance and growth

We first tested how the MDR plasmids pLL35 and pOXA-48 affected susceptibility to antibiotics. To do this, we determined MIC curves across concentration gradients of CEF and ERT in transconjugants harbouring pLL35 or pOXA-48 or both plasmids, alongside plasmid-free controls (Fig. 1). Each plasmid provided a distinct resistance profile, combining to provide high-level resistance to both antibiotics in strains carrying both plasmids (Two-way ANOVA Interaction F3,46 = 218.1, p < 0.01). Acquiring pLL35 provided high level resistance to CEF (MIC = >128 μg/ml) without cross-resistance to ERT (MIC = 0.05 μg/ml). In contrast, acquiring pOXA-48 provided variable resistance to ERT (MIC range 6 - 24 μg/ml) and moderate cross-resistance to CEF (MIC range 8 - 64 μg/ml). However, there was no correlation observed between levels of resistance to ERT and CEF among the pOXA-48 transconjugants, indicating that the level of resistance to ERT does not predict the degree of cross-resistance to CEF (Spearman’s rank correlation, ρ = 0, p-value = 1). Acquiring both pLL35 and pOXA-48 provided resistance to both ERT and CEF (ERT MIC range 3-12 μg/ml; CEF MIC = >128 μg/ml).

**Figure 1:**
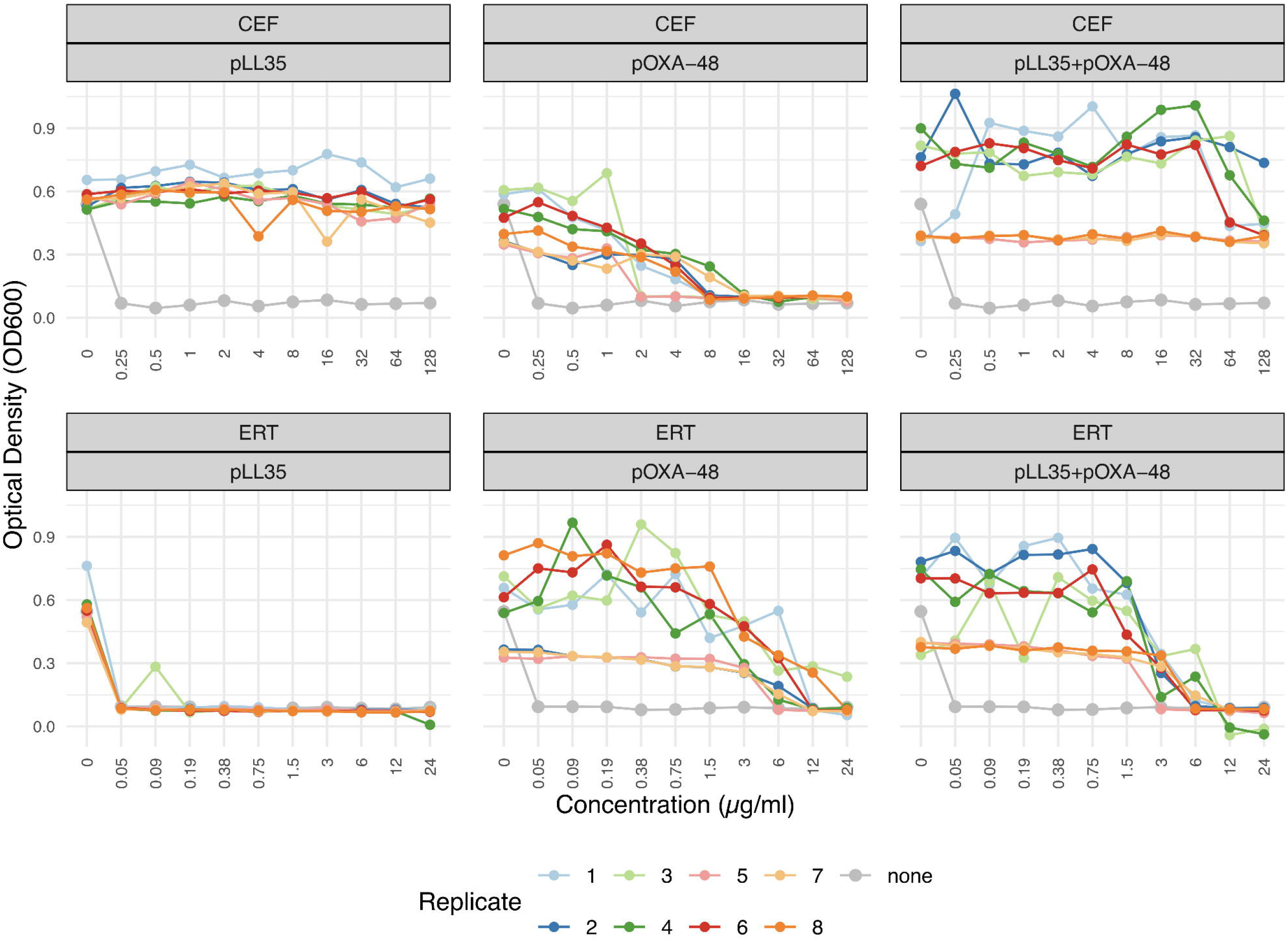
Resistance profiles of *E. coli* MG1655 isolates harbouring pLL35, pOXA-48 or both. MIC curves show optical density (OD600) measurements for pLL35, pOXA-48, and pLL35+pOXA-48 transconjugants in the presence of increasing concentrations of cefotaxime (CEF, top panels) and ertapenem (ERT, bottom panels). Each point represents mean of 3 technical replicates for each of the eight biological replicates (colored lines 1-8); plasmid-free ancestral *E. coli* MG1655 control is shown in grey.

We observed high variability in optical density within MICs among transconjugants with pOXA-48 or both plasmids, but not when harbouring pLL35 alone. This suggested that the acquisition of pOXA-48 differentially impacted growth of each transconjugant. To investigate these growth effects further, we obtained 24-hour growth curves for all strains (Fig. 2). Transconjugants carrying pOXA-48 or both pOXA-48 plus pLL35 showed greater variance in multiple growth kinetic parameters compared to those carrying only pLL35 (MaxGrowthRate: pLL35 vs pOXA_48: F(13,15) = 0.133, p = 0.00075; pLL35 vs pLL35 + pOXA_48: F(13,23) = 0.143, p = 0.00075; Integral: pLL35 vs pOXA_48: F(13,15) = 0.016, p < 0.00001; pLL35 vs pLL35 + pOXA_48: F(13,23) = 0.029, p < 0.00001; LagTime: pLL35 vs pOXA_48: F(13,15) = 0.195, p = 0.00523; pLL35 vs pLL35 + pOXA_48: F(13,23) = 0.314, p = 0.034). Moreover, pOXA-48 and pLL35 differed in their impact upon the growth kinetics of *E. coli* (ANOVA: F3,56 = 6.647, p = 0.000641), which was primarily driven by pOXA-48 reducing maximum growth rate (diff = -0.01644, p = 0.0096 vs. none), maximum growth (max OD: diff = - 0.03668, p = 0.6794 vs. none), and the area under the growth curve (Integral: diff = - 1.31127, p = 0.0708 vs. none). Together, these data suggest that pOXA-48 is costlier than pLL35, but that the effects of pOXA-48 on bacterial growth vary markedly between individual transconjugants.

**Figure 2:**
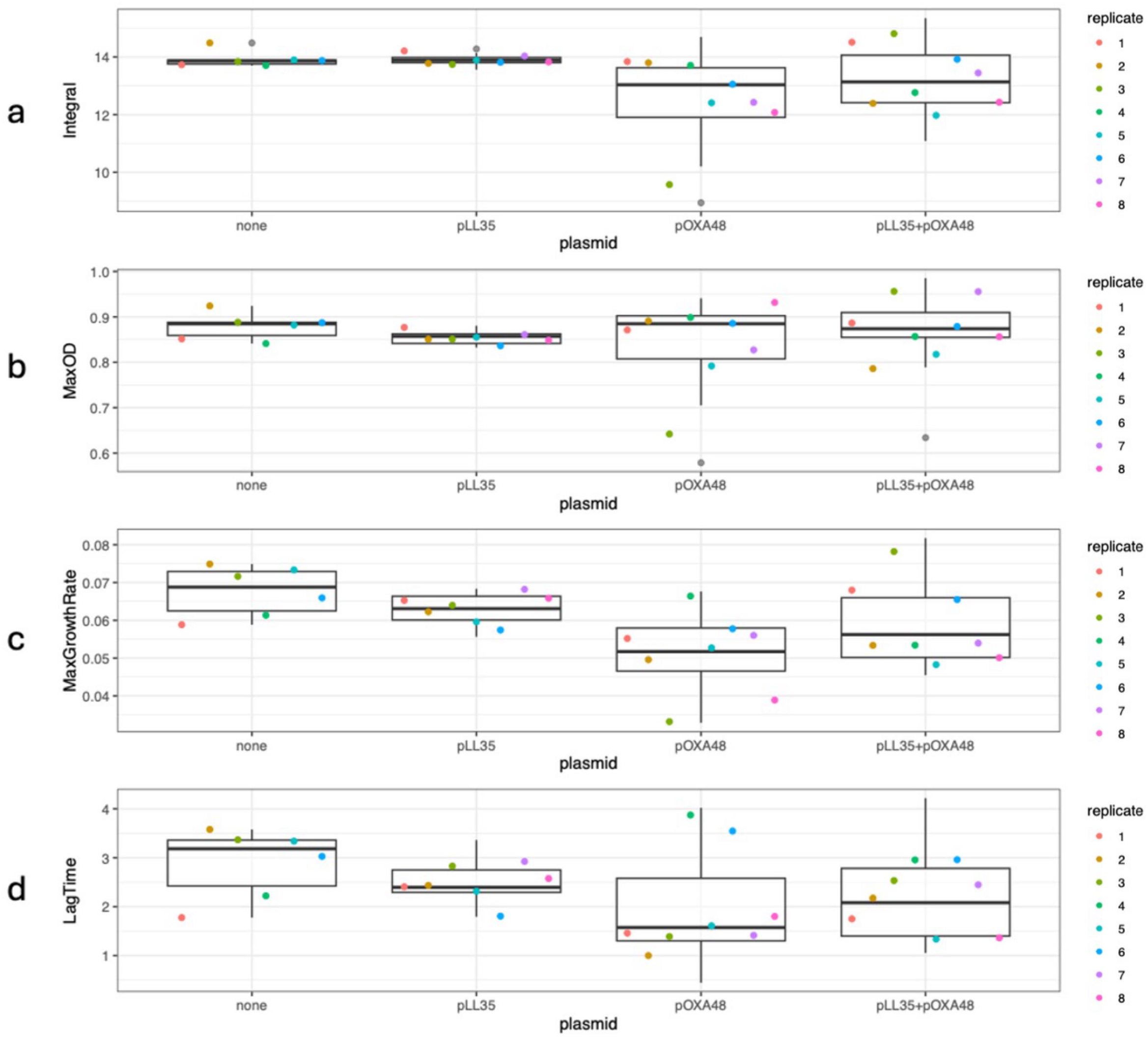
The effect of the acquisition of pOXA-48, pLL35 or both plasmids on the growth kinetics of *E. coli* transconjugants. Each coloured point represents the mean of 3 technical replicates for each biological replicate (n = 8). Horizontal lines within the box plots represent median values, the upper and lower hinges are the 25^th^ and 75^th^ percentiles and the whiskers extend to observations within 1.5 X the interquartile range. Panels represents the effect of plasmid(s) acquisition on **a**, integral (area under the growth curve) **b**, max OD **c**, max growth rate **d**, and lag time for transconjugants with different plasmids, in addition to the plasmid-free control (labelled none).

### Mutations obtained by transconjugants during plasmid acquisition

We hypothesised that the variability in growth kinetics caused by pOXA-48 could be due to differential mutations gained within each transconjugant during plasmid acquisition. To test this, sequenced the genomes of each independent transconjugant. 15 of 24 transconjugants had at least one additional chromosomal mutation, whilst none had mutations in either plasmid (Fig. 3). Whereas 8/8 transconjugants with both plasmids and 6/8 transconjugants with pOXA-48 alone had gained additional mutation(s), only 1/8 transconjugants with pLL35 did (Table S3). By far the most commonly mutated gene was *ompF*, encoding an outer membrane pore-forming protein associated with antibiotic permeability [22]. Mutations were detected in five transconjugants with both plasmids—four resulting in frameshifts and one in a premature stop codon—and in one transconjugant carrying only pOXA-48, resulting in a frameshift. Notably, three additional mutations were observed in other transconjugants carrying either pOXA-48 or both plasmids that are likely to affect expression of *ompF*. These included both genes of the *envZ-ompR* two-component system regulating *ompF* having missense mutations [30], and an intergenic mutation located 50 bp from the ompF transcription start site. In total, 9/16 transconjugants containing pOXA-48 had gained additional mutations affecting *ompF* or its regulation.

**Figure 3:**
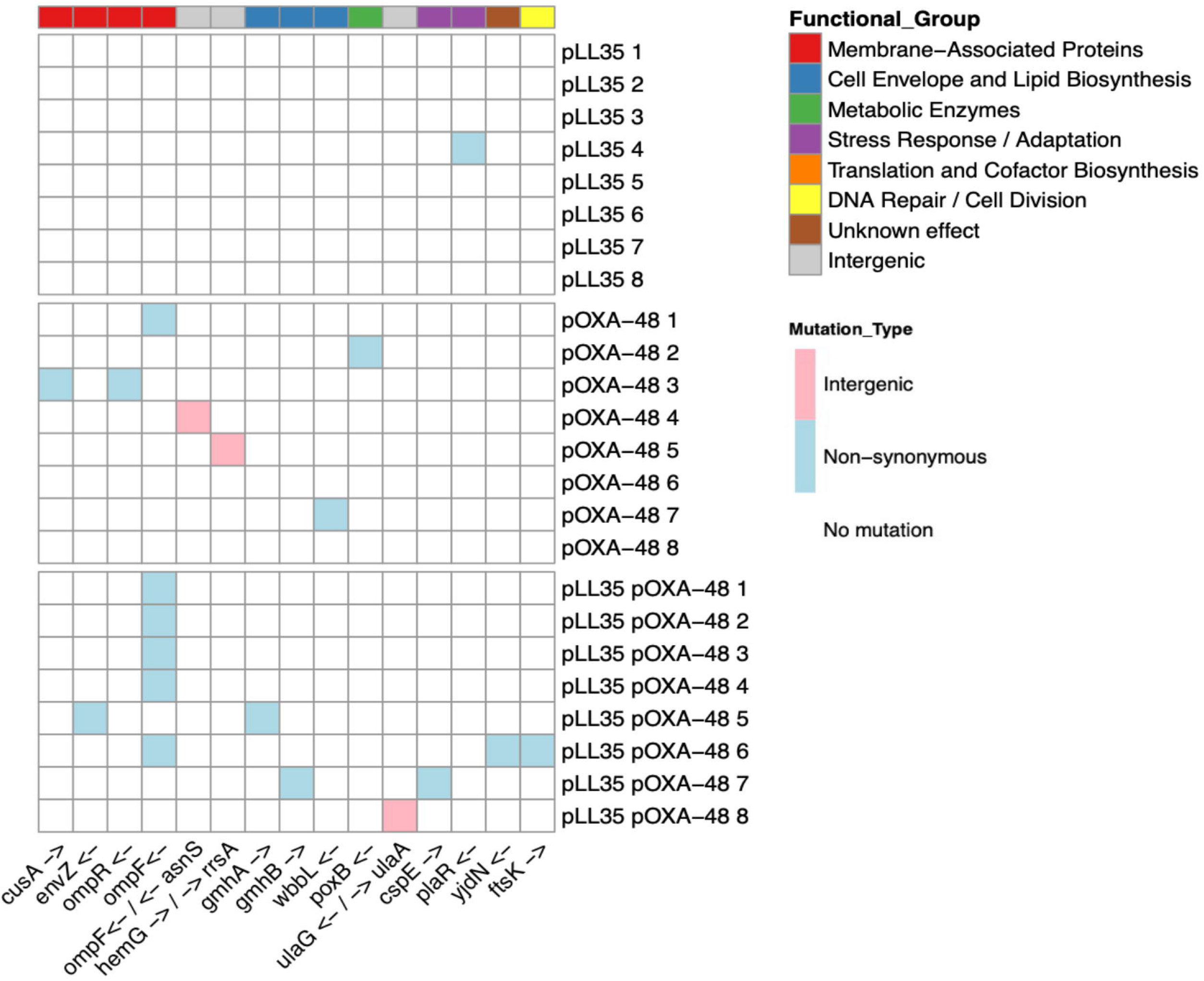
Heatmap of the distribution of chromosomal mutations across replicate *E. coli* transconjugants harbouring either pLL35, pOXA-48, or both plasmids. Each row corresponds to an independent transconjugant, and each column corresponds to a mutation locus. Pink, intergenic mutations, blue represents non-synonymous mutations and white no variants observed. Loci are clustered into functional groups denoted by the colours at the top of the plot with grey boxes representing intergenic mutations.

Given the well-established role of OmpF in cellular antibiotic permeability [22, 30, 31], we next tested how loss of *ompF, ompR* or *envZ* altered ERT resistance with or without pOXA-48 using single gene knockout strains for each gene from the Keio library (Fig. 4). Whereas none of these mutations increased ERT resistance without the plasmid, each was strongly synergistic with pOXA-48, amplifying the ERT resistance level provided by the plasmid with the strongest effect seen for loss of *ompR* (Fig. 4, ANOVA AUC, F3,16=7.499, P=0.0024; Post-hoc Tukey Tests, *ompR* pOXA_4:envZ pOXA_48, P = 0.00595, *ompR* pOXA_48:*ompF* pOXA_48, P = 0.01578). Next, using 24-hour growth curves we tested the effect of these chromosomal mutations on pOXA-48 fitness costs (Fig. 5).

**Figure 4:**
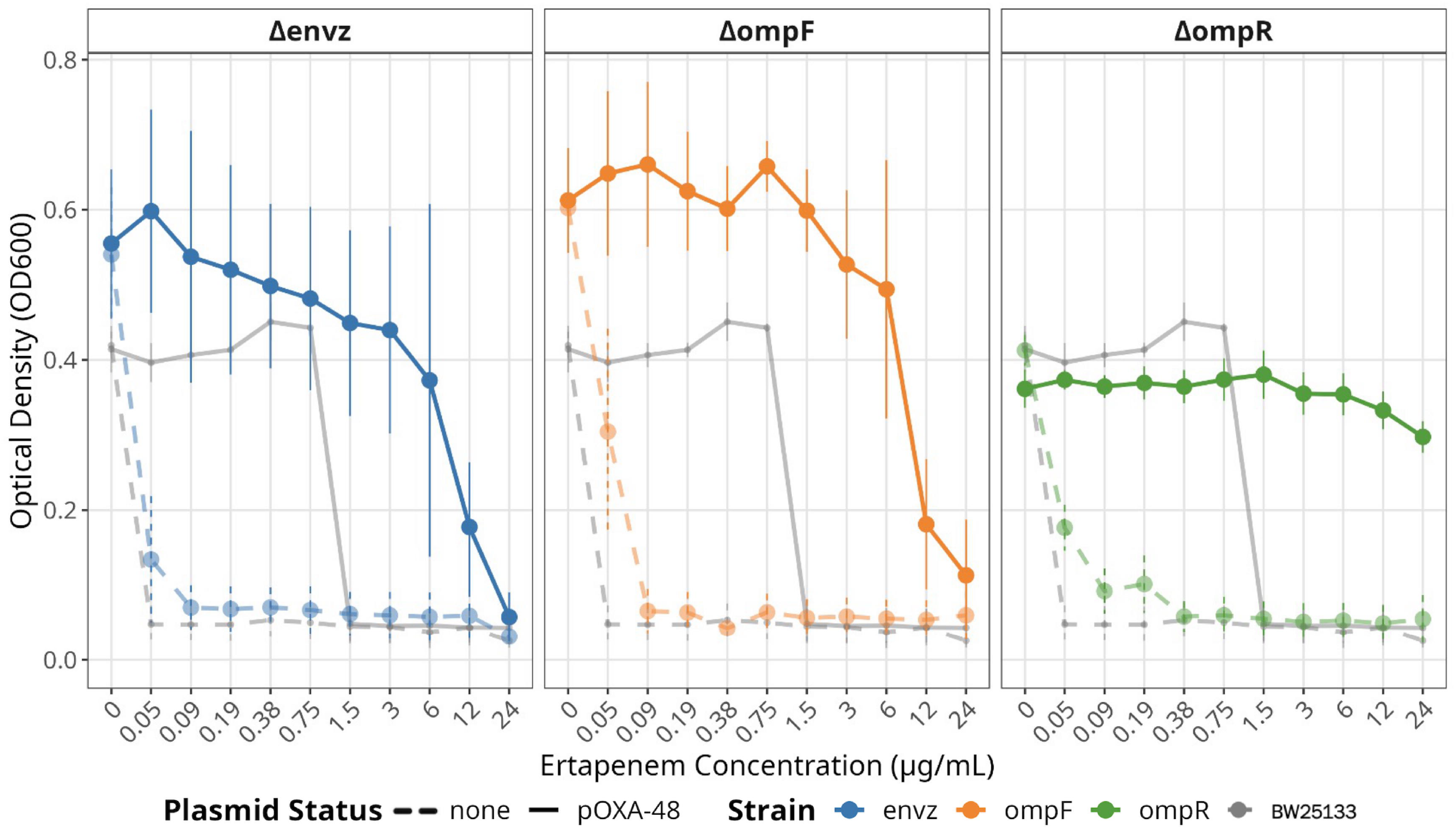
Ertapenem resistance profiles of *E. coli* BW25113 and knockout mutants with and without the pOXA-48 plasmid. Each point represents the mean growth (OD600) of three replicates per strain at each drug concentration, in the presence of increasing ertapenem concentration. Plot is faceted by gene knockout; dashed lines show growth with no plasmid and solid lines show growth harbouring pOXA-48. Grey lines in each panel show the growth of WT *E. coli* BW25113 for reference.

**Figure 5:**
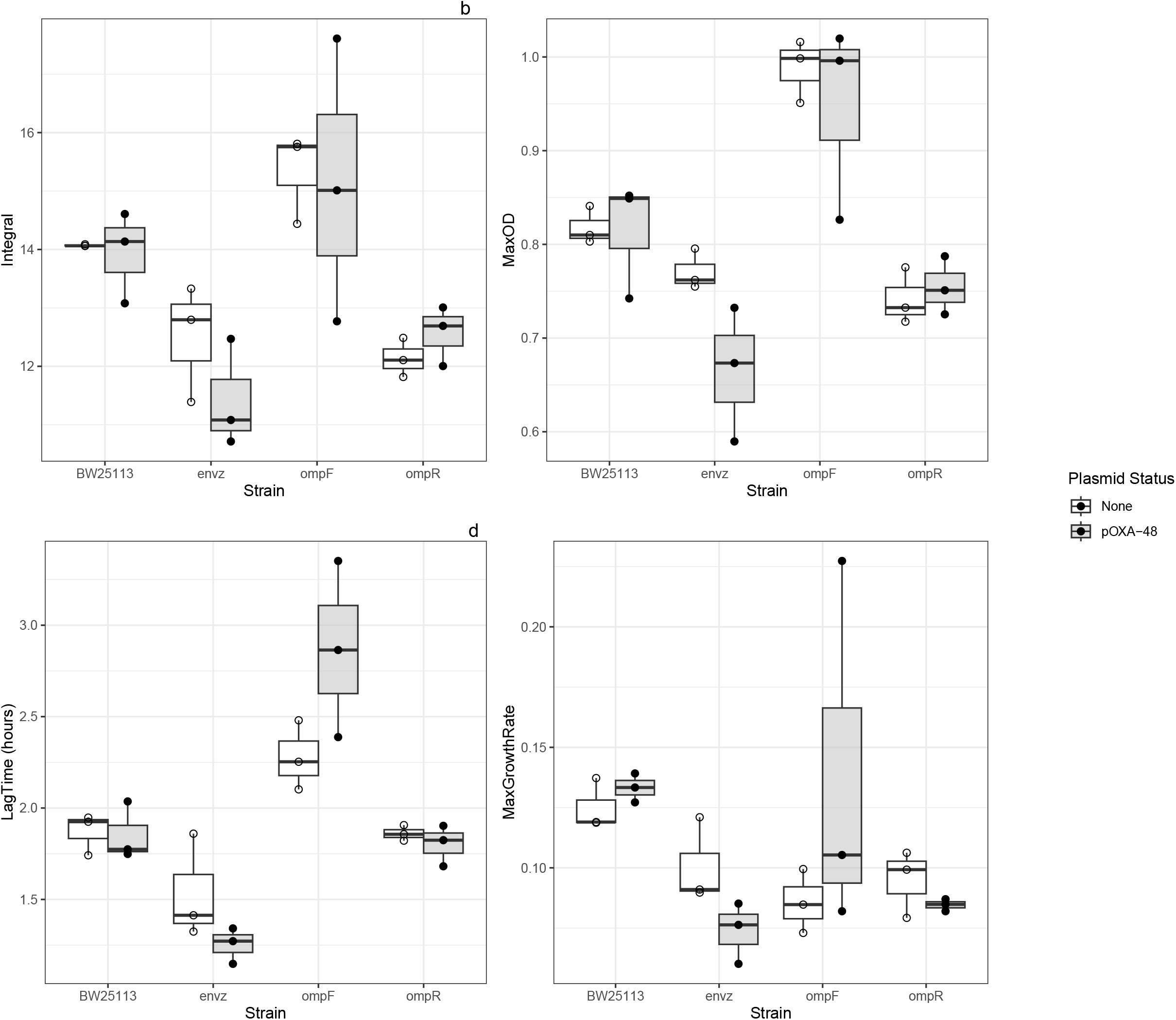
The effect of the acquisition of pOXA-48 on the growth kinetics of the knockout strains; *ΔenvZ, ΔompF* and *ΔompR* and WT *E. coli* BW25113. Grey-filled boxplots represent plasmid-bearing strains, and white (unfilled) boxplots represent their plasmid-free counterparts. Each point represents the mean of 3 technical replicates for each of 3 biological replicates. Horizontal lines within the box plots indicate median values; the upper and lower hinges are the 25^th^ and 75^th^ percentiles and the whiskers extend to observations within 1.5× the interquartile range. Panels a– d show the effect of pOXA-48 acquisition on growth parameters by comparing plasmid-free and plasmid-bearing versions of knockout strains: a – integral (area under the curve), b – maximum OD_600_, c – lag time, and d – maximum growth rate.

We observed no compensatory effect of losing any of these genes on the growth of plasmid carriers (ANOVA max growth rate, F1,52=0.511, P=0.47805), indeed loss of *envZ* strongly impaired growth both with and without pOXA-48 (Post-hoc Tukey Tests, envZ:BW25113, P=0.0013; envZ pOXA-48:BW25113 pOXA-48, P=0.0046), whereas loss of *ompF* increased growth regardless of plasmid carriage (Post-hoc Tukey Tests, ompF pOXA-48:ompF none, P=0.0490). These effects are consistent with the specific mutations observed in the transconjugants - the frameshift and nonsense mutations in *ompF* likely results in loss of functions similar to our knockout strains, while the missense mutation in envz (I86S) may have more subtle effects than total deletions (Table S3). Together these patterns suggest that mutation of *ompF* and its regulators in transconjugants was primarily favoured due to the synergistic effects on ERT resistance, rather than compensatory effects on fitness.

## Discussion

We show that additional chromosomal resistance mutations occurring during the formation (or outgrowth) of transconjugant colonies facilitated the acquisition of pOXA-48 but not during the acquisition of pLL35. These mutations affected the outer membrane protein OmpF, or its regulators *ompR* or *envZ*. OmpF is known to control antibiotic permeability, including of carbapenems [30, 32]. Loss of *ompF* or either regulator did not alone substantially increase ERT resistance but instead these chromosomal mutations acted synergistically with the OXA-48 ARG encoded by pOXA-48 to greatly increase ERT resistance in plasmid carriers. None of these mutations were compensatory for the plasmid, suggesting that it was their effect on resistance level rather than any compensatory effect upon fitness that led to their enrichment in pOXA-48 transconjugants. Overall, our results highlight how chromosomal resistance and acquired ARGs can have strong epistatic effects on resistance level, here acting synergistically to provide greater than additive levels of resistance to ERT and facilitating the acquisition of MDR plasmids in antibiotic environments.

Previous studies report synergy between chromosomal and plasmid encoded resistance. Through a synergistic pairing of reduced tetracycline influx, via the loss of function of *ompF*, and efflux via a plasmid encoded tetracycline efflux pump, *E. coli* can modulate resistance; increasing the level of resistance while minimising costs [19]. Synergy between antibiotic inactivation and influx has also been reported; the trace catalytic activity of the plasmid encoded ESBL CTX-M-15 and reduced drug influx has previously been shown to promote the evolution of carbapenem resistance through the altered expression of porins leading to increase resistance to ERT [30] Coupling of β-lactamase production and porin inactivation has also been observed within non-carbapenemase-producing CRE isolates from patients with bacteraemia, resulting in high level carbapenem resistance [33]. Similarly, transconjugants receiving plasmids encoding New Delhi metallo-β-lactamase 1 (*bla*_NDM-1_) rapidly acquired mutations in *ompR* altering the expression of *ompC* and *ompF*, when selected for using the carbapenem meropenem [34]. In our case, synergy arises between a plasmid encoded carbapemenase OXA-48, which alone provides resistance to ERT, and mutations which act to lower antibiotic permeability. Together these are likely to reduce the rate of periplasmic accumulation of antibiotic [35], resulting in a reduction of the enzymatic activity required to overcome lethal concentrations of carbapenems [6], thus providing high level resistance.

Rapid compensatory evolution during transconjugant formation can stabilise plasmids [21]. Here, we demonstrate another way in which rapid evolutionary dynamics can facilitate plasmid-mediated HGT, specifically by chromosomal resistance mutations enhancing the fitness benefits of the plasmid encoded ARGs under antibiotic selection. Such rapid evolutionary processes may be important for understanding plasmid dynamics in environments with strong antibiotic selection for plasmid-encoded ARGs, and especially where these ARGs provide only marginal resistance, as here with ERT and pOXA-48. In contrast, when plasmid-encoded resistance mechanisms provide very high-level resistance, such as CTX-M-15 with CEF, no additional resistance mechanisms were required for the stable plasmid transfer during CEF exposure. Additional chromosomal mutations were also required during the acquiring both pLL35 and pOXA-48 under multidrug treatment, indicating that there was no synergy between the two plasmids in resistance to ERT.

Our findings are consistent with clinical observations, where pOXA–48–producing Enterobacterales frequently exhibit mutations in genes encoding outer membrane protein or their regulators, such as *ompR* and *ompF*, that reduce outer membrane permeability [30, 31]. For instance, clinical isolates of *Klebsiella pneumoniae* with alterations in *ompC* and *ompF* demonstrate significantly reduced susceptibility to carbapenems, including ertapenem and meropenem [22]. These chromosomal resistance mutations, together with plasmid-borne beta-lactamases, create highly resistant clinical strains, reducing treatment options. Moreover, synergistic epistasis between chromosomal and plasmid encoded resistance determinants is likely to complicate the prediction of antibiotic resistance level from genomic data.

## Funding

KNH is supported by the Petroleum technology Development Fund (PTDF) Nigeria Scholarship (PTDF/ED/OSS/PHD/KNH/1681/2019PHD263). MJB is supported by the Wellcome Trust (221653/Z/20/Z).

## Acknowledgement

We would like to thank Rok Krašovec for providing the Keio knockout strains, Álvaro San Millán for the *E. coli* J53 pOXA-48 strain and, and Laura Carrilero for sharing the pLL35 plasmid. We also thank Steven Dunn for providing the sequence annotation for the pLL35 plasmid.

## Author contributions

Conceptualization MAB, MJB, Data curation KNH, Formal analysis KNH, Funding acquisition KNH, MAB, MJB, Investigation KNH, Methodology MAB, MJB, Project administration MAB, MJB, Resources MAB, MJB, Supervision MAB, MJB, Visualization KNH, Writing – original draft KNH, Writing – review & editing MAB, MJB.

## Conflict of Interest

The authors declare that there are no conflicts of interest.

## Notes

### Competing Interest Statement

The authors have declared no competing interest.

