## Supplementary Figures for "Chromosomal resistance mutations facilitate acquisition of multidrug-resistance plasmids in *Escherichia coli*"

### SUPPLEMENTARY MATERIAL_FIGURES

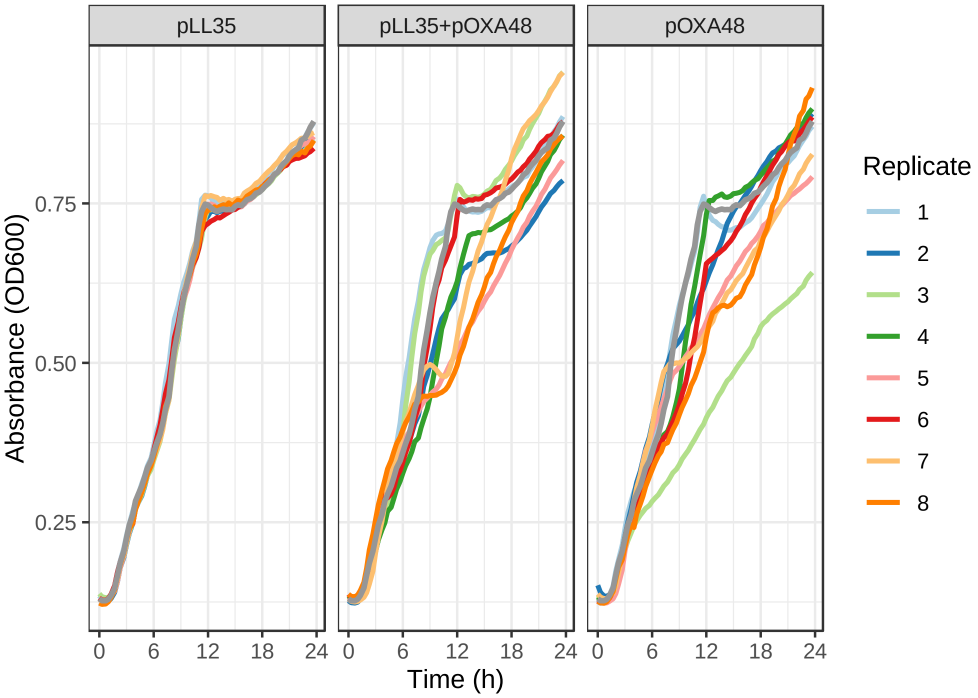


Figure S1: Growth curve of 8 transconjugants per plasmid type with OD600 plotted against time in hours. The grey line overlayed is the reference plasmid-free ancestral plasmid. Each of the coloured lines is the average of 3 replicates of each individual replicate in the three plasmid groups.

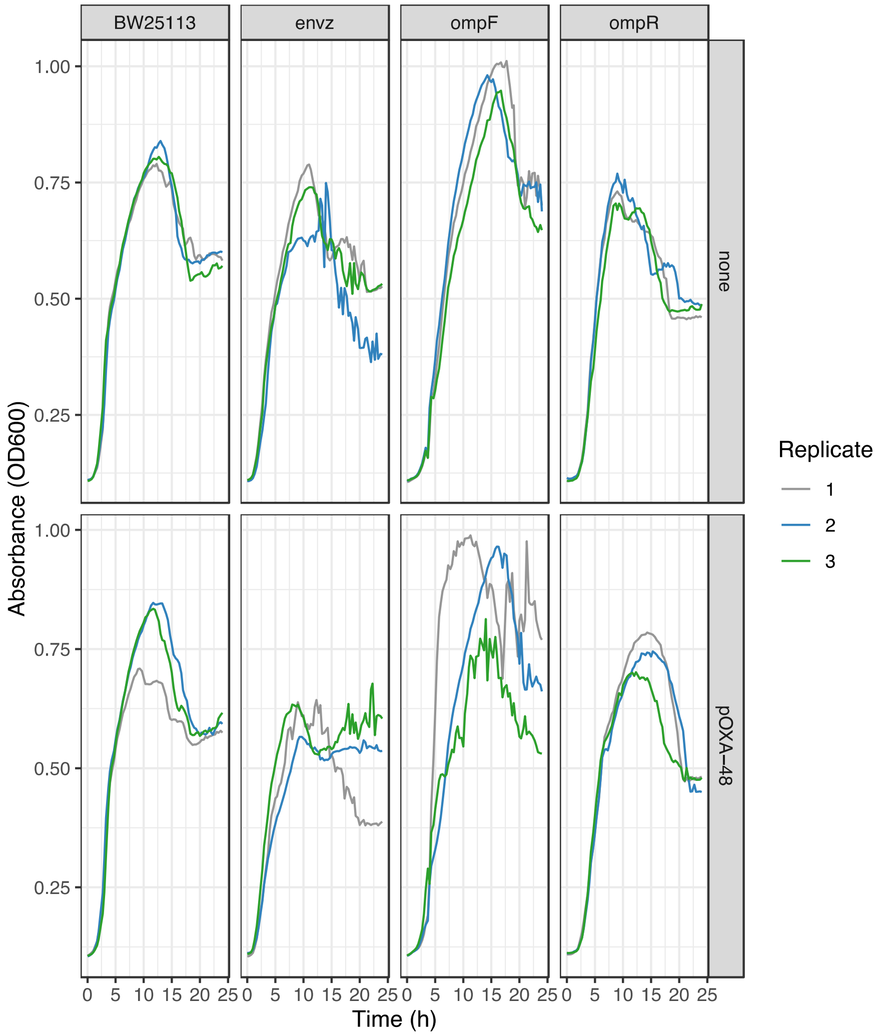


Figure S2: Growth Curves of Keio Knockouts and BW25113 Strains with and without pOXA-48 Plasmid. Figure shows the 24-hour growth curves (OD600 absorbance) of E. coli BW25113 and Keio knockout strains (*envZ*, *ompF*, *ompR*) with and without the pOXA-48 plasmid. The top panel presents growth curves for plasmid-free strains, and the bottom panel shows strains carrying the pOXA-48 plasmid. Each plot represents a specific strain, with each line being the mean of three 3 technical replicates. The Y-axis represents the optical density at 600 nm (OD600), which serves as a measure of cell growth, and the X-axis represents the growth time in hours.
